# A *Legionella* effector ADP-ribosyltransferase inactivates glutamate dehydrogenase

**DOI:** 10.1101/2020.08.03.234963

**Authors:** Miles H. Black, Adam Osinski, Marcin Gradowski, Kelly A. Servage, Krzysztof Pawłowski, Vincent S. Tagliabracci

## Abstract

ADP-ribosyltransferases (ARTs) are a widespread superfamily of enzymes frequently employed in pathogenic strategies of bacteria. *Legionella pneumophila*, the causative agent of Legionnaire’s disease, has acquired over 330 translocated effectors that showcase remarkable biochemical and structural diversity. Here we took a bioinformatic approach to search the *Legionella* effector repertoire for additional divergent members of the ART superfamily and identified an ART domain in Lpg0181. We show that *L*. *pneumophila* Lpg0181 targets a specific class of 120-kDa NAD+-dependent glutamate dehydrogenase (GDH) enzymes found in fungi and protists, including many natural hosts of *Legionella*. Lpg0181 targets a conserved arginine residue in the NAD+ -binding pocket of GDH, thereby blocking oxidative deamination of glutamate. While intracellular pathogens employ diverse virulence mechanisms to overcome host-limited nutrient availability, Lpg0181 is––to the best of our knowledge––the first example of a *Legionella* effector which directly targets a host metabolic enzyme.

## Introduction

Our lab has previously taken a bioinformatic approach to identify atypical and uncharacterized members of the protein kinase superfamily. Several of the outlying kinase families we have characterized are components of bacterial conflict systems, including the HopBF1 substrate of the *Pseudomonas syringae* type III secretion system (1) and the Lpg2603 and SidJ substrates of the *Legionella pneumophila* Type IV secretion system (T4SS) (2,3). *Legionella* is a gram-negative environmental pathogen and the causative agent of a potentially fatal pneumonia called Legionnaire’s disease. The genus *Legionella*, through horizontal gene transfer from its hosts and cohabiting bacteria, has acquired over 18,000 translocated effectors representing at least 137 different eukaryotic-like domains (4). *Legionella pneumophila* alone translocates more than 330 effectors, accounting for about 10% of its proteome (5). Because these effectors have evolved to target conserved processes, they represent an orthogonal approach to interrogate eukaryotic biology. Furthermore, effectors are a rich source of structural and biochemical diversity; of the 99 conserved protein domains identified in *L*. *pneumophila* effectors, 46 are entirely novel (5). Even those effectors with recognizable protein folds sometimes catalyze unexpected reactions (6-8). For instance, we recently characterized the SidJ effector ‘pseudokinase’, which adopts a protein kinase fold, but catalyzes polyglutamylation (3,9-11). SidJ inactivates the SidE effectors, which harbor phosphodiesterase (PDE) and ADP-ribosyltranferase (ART) domains that cooperate to catalyze phosphoribosyl-linked ubiquitination independent of host E1 and E2 enzymes (12-19).

Inspired by the SidE effectors, we sought to identify novel ART folds in the *Legionella* effector repertoire. Members of the ART superfamily transfer ADP-Ribose (ADPR) from NAD+, joining the 1′ position of the ribose in N-, S-, or O-linked glycosidic bonds to diverse substrates including proteins, nucleic acids, and small molecules (20). ARTs are widespread in nature, but have extensively diversified in conflict-related systems from bacteria (21). While poly-ADP-ribosylation is found almost exclusively in multicellular eukaryotes, the prokaryotic ARTs transfer a single ADPR moiety to their substrates (22). Mono-ADP-ribosylation by bacterial toxins, including diphtheria, cholera, pertussis and clostridial toxins, is a fundamental pathogenic mechanism for many serious human diseases (23). While the conventional function of ADP-ribosylation is to lock host proteins in a permanently active or inactive state, the SidE ART domain from *Legionella* reveals that ADP-ribosylation can be an intermediate for other types of unique chemistry. SidE is the only ART domain identified thus far among *Legionella* T4SS effectors. Here we have discovered a mono-ART fold in the *Legionella* effector Lpg0181, which bears only 19% sequence identity with the SidE ART domain. Lpg0181 was identified as a substrate of the *L*. *pneumophil*a T4SS in a screen based on translocation of a β-lactamase fusion (24), but its activity and substrates are unknown.

## Results

### Identification of a mono-ADP-ribosyltransferase fold in the Lpg0181 family of Legionella effectors

We applied a bioinformatic strategy to search for outlying members of the ADP-ribosyltransferase (ART) superfamily (**Fig. S1**).

Using the FFAS sequence profile algorithm (25), we identified remote similarity to ARTs in the Lpg0181 family of proteins from the human pathogen *Legionella*. Its closest relatives by FFAS include butterfly DNA ADP-ribosyltransferase pierisin, *Salmonella* typhoid toxin, and pertussis toxin with 12-16% sequence identity and significant FFAS Z-scores between - 10 and -11. The predicted ART domain of the *L*. *pneumophila* Lpg0181 lies between residues 1 and 174, while residues 174-304 are predicted to adopt a coiled-coil structure (**Fig. 1*A***). Sequence alignments identified three conserved active site motifs (R37, ^86^SxS and ^135^ExE), which place the Lpg0181 family in the R-S-E clade of mono-ARTs (mARTs) (21,26) and suggest it should be enzymatically active (**Fig. 1*B***). The lpg0181-centered family is found only in the genera *Legionella* and *Fluoribacter*.

**Figure 1.**
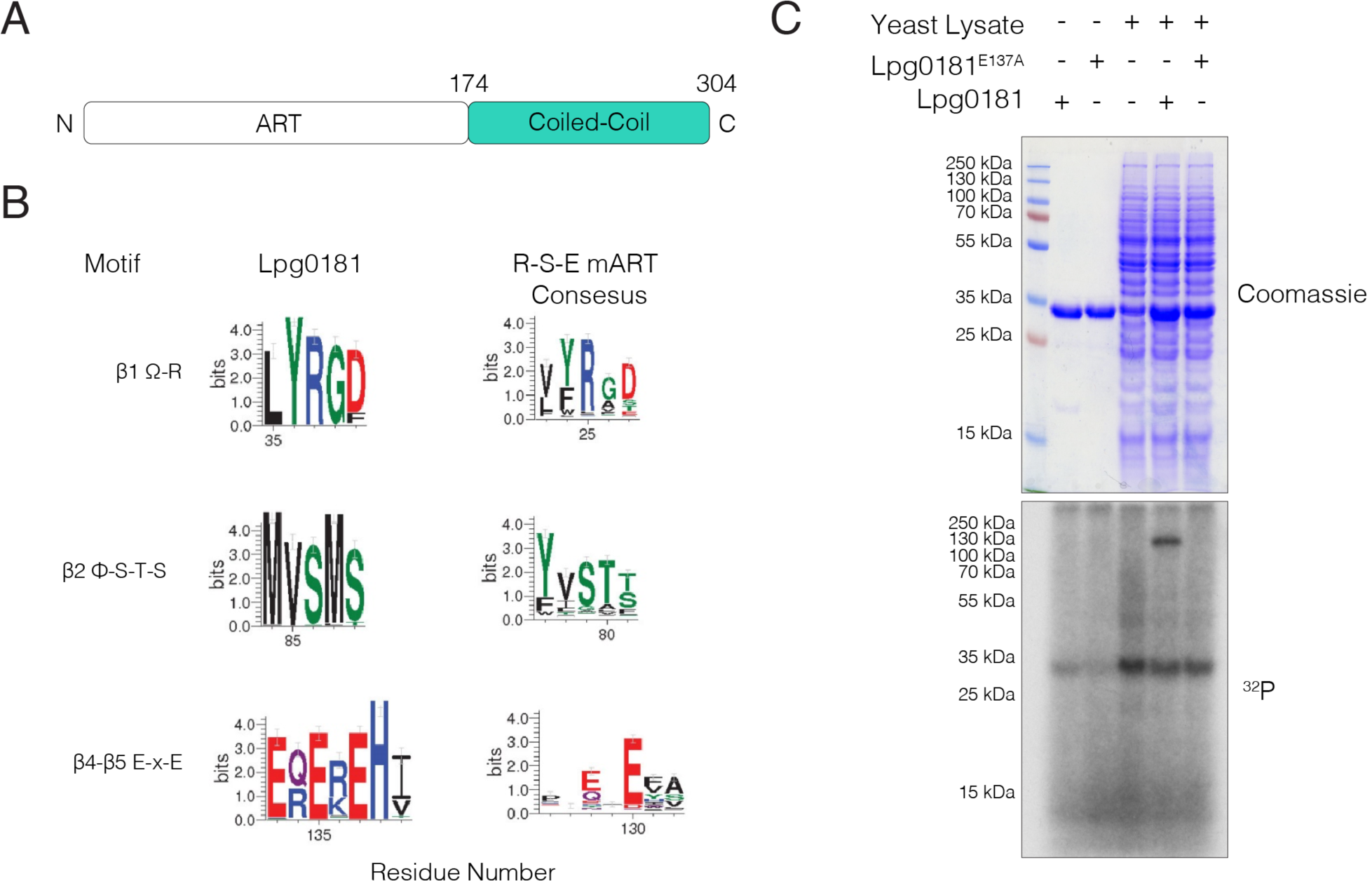
Lpg0181 is a mono-ADP-ribosyltransferase that targets a ∼120 kDa protein in yeast. **(A)** Domain schematic of Lpg0181. **(B)** Sequence logos (weblogos) illustrating the conservation of predicted catalytic residues (26) in 81 Lpg0181 homologs (left) and 453 members of the R-S-ExE clade of mARTs (pfam PF01375, enterotoxin a) (right). **(C)** Incorporation of [^32^P] from [^32^P]-adenylate NAD+ into an unknown 120 kDa band in a yeast lysate by Lpg0181, but not the E137A mutant.

### Lpg0181 ADP-ribosylates yeast glutamate dehydrogenase 2 (Gdh2)

To identify substrates of this putative *Legionella* ART, we incubated purified recombinant Lpg0181 with yeast cell extracts and [^32^P]-adenylate NAD+. Lpg0181, but not the E137A mutant, incorporated ^32^P into a species in a yeast extract that migrated at ∼120 kDa during SDS-PAGE (**Fig. 1*C***). There were no substrates labeled by Lpg0181 when mammalian cell lysates were tested (not shown). To identify the ∼120 kDa substrate, we repeated the labeling experiment with biotin-17-NAD+ and used streptavidin resin to enrich biotin-labeled proteins. Bound proteins were eluted by trypsinization and identified by LC-MS/MS. We prioritized candidate substrates by only considering proteins enriched from reactions containing Lpg0181 but not the E137A mutant. This step eliminated the most abundant proteins identified in both streptavidin pulldowns, including several carboxylase enzymes, which are covalently modified by a biotin prosthetic group. The list of unique proteins was then sorted by MASCOT protein score. The top hit from this sorting was NAD+-dependent glutamate dehydrogenase 2 (Gdh2), calculated molecular weight of 124.3 kDa) (**Table S1**).

### Lpg0181 specifically modifies fungal and protozoan isoforms of glutamate dehydrogenase

GDH enzymes can be broadly divided into four classes *(27,28)*. The two most common and by far most-studied GDH classes are small (∼50 kDa), hexameric enzymes with widespread phylogenetic distributions. These include the vertebrate GDH enzymes, which are extensively regulated by allosteric feedback (29). A third class of ∼180 kDa, NAD^+^-dependent GDH is found only in eubacteria. A fourth class of GDH comprises a group of tetrameric, ∼115 kDa enzymes found in fungi, protists, and some deltaproteobacteria (**Fig. 2*A***). yeast Gdh2 belongs to the 115 kDa-class, catalyzes glutamate deamination, and is required for normal yeast growth with glutamate as a sole nitrogen source (30,31). Although yeast also possess two GDH enzymes of the widespread 50-kDa class (32,33), neither were identified in our streptavidin pulldowns. Notably, the 115-kDa class is also present in protozoa, the natural hosts of *Legionella*. We selected representative GDH enzymes for heterologous expression in *E*. *coli* and tested whether recombinant GDH are substrates of Lpg0181 in reactions containing [^32^P]-NAD+. Lpg0181 ADP-ribosylated yeast Gdh2 and Glud2 from the amoeba *D*. *discoideum (Dd*Glud2) but did not ADP-ribosylate human or *Legionella* GDH homologs, nor did it modify yeast Gdh1 (**Fig. 2*B***). Thus, of the substrates we tested, only GDH enzymes of the 115-kDa class are recognized by Lpg0181.

**Figure 2.**
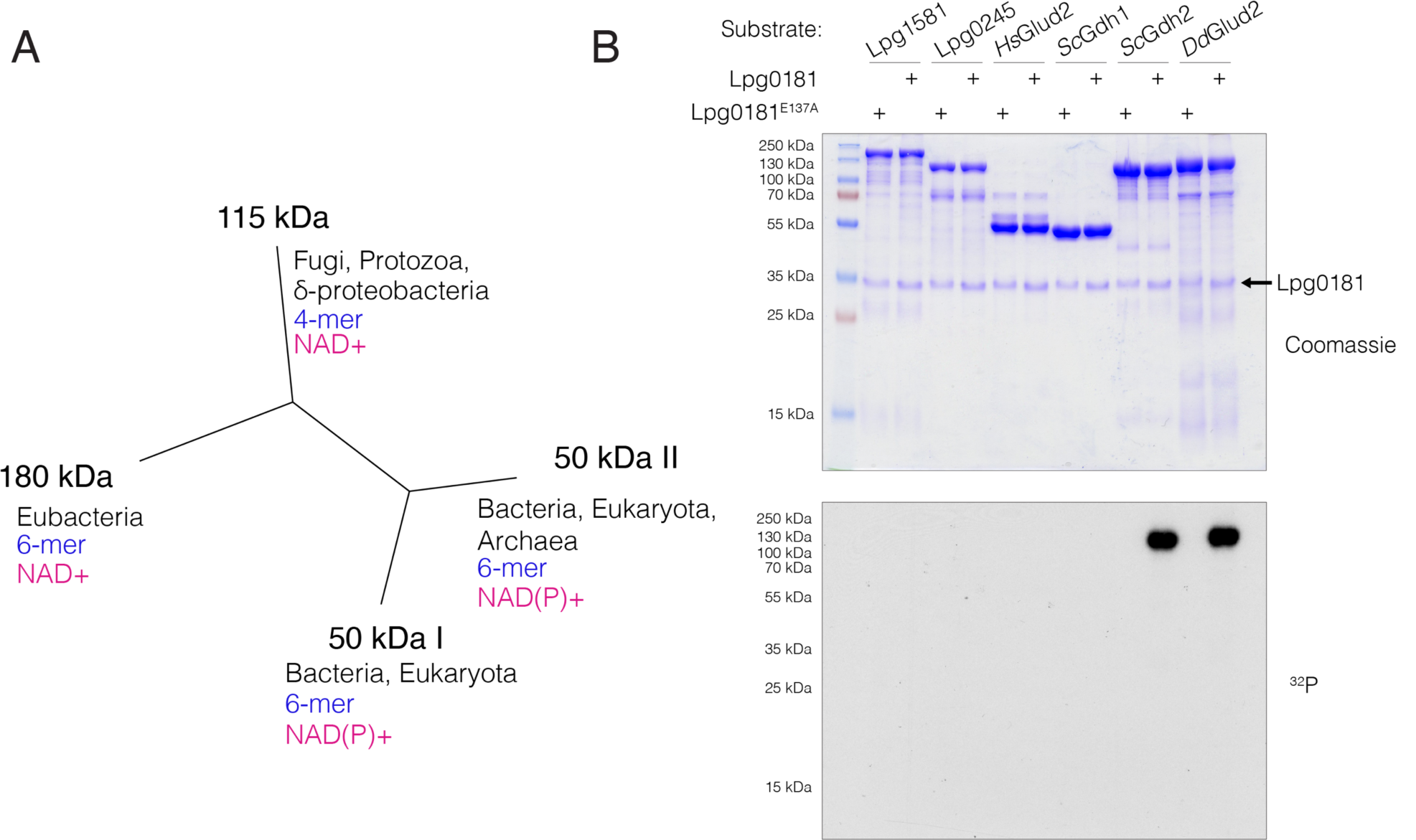
Lpg0181 ADP-ribosylates fungal and protozoan isoforms of glutamate dehydrogenase. **(A)** Dendrogram depicting the four classes of glutamate dehydrogenase isoforms labeled by their subunit molecular weight, adapted from Miñambres et. al (27). Each class is annotated with its phylogenetic distribution (black), monomer composition (blue), and cofactor preference (magenta). **(B)** Incorporation of [^32^P] from [^32^P]-NAD+ by Lpg0181 or the inactive E137A mutant with a panel of glutamate dehydrogenases as substrates: the two *Legionella* glutamate dehydrogenases (Lpg1581 and Lpg0245), human Glud2 (*Hs*Glud2), yeast (*Sc*Gdh1 and *Sc*Gdh2), and *Dictyostelium* Glud2 (*Dd*Glud2). Reaction products were separated by SDS-PAGE and incorporated radioactivity visualized by autoradiography.

Lpg0181-dependent ADP ribosylation of *Dd*Glud2 was time-dependent (**Fig. S2*A***) and displayed a pH optimum of 6.0 and a slight preference for low (0-50 mM) salt (**Fig. S2*B* and S2*C***). *Dd*Glud2 was not ADP-ribosylated in reactions without ART enzyme or with ART domains from other *Legionella* effectors, and Lpg0181-catalyzed incorporation of etheno-ADP-Ribose occurred even in the presence of excess unlabeled ADP-ribose (**Fig. S3**). We determined the kinetic parameters of Lpg0181 with *Dd*Glud2 as a substrate. The Km for *Dd*Glud2 was 2.22 *μ*M (95% CI 1.54, 3.16) and the Km for NAD+ was 1.57 *μ*M (95% CI 1.26, 1.95) (**Fig. 3*A* and 3*B***), in accordance with eukaryotic cytosol NAD+ concentrations of ∼0.3 mM (34). Lpg0181 has a turnover rate (K_cat_) of 0.15 min^-1^. After a complete (1.5 h) reaction, intact mass analysis of *Dd*Glud2 revealed compete stoichiometric addition of a single ADP Ribose moiety (+541 Da) (**Fig. 3*C***).

**Figure 3.**
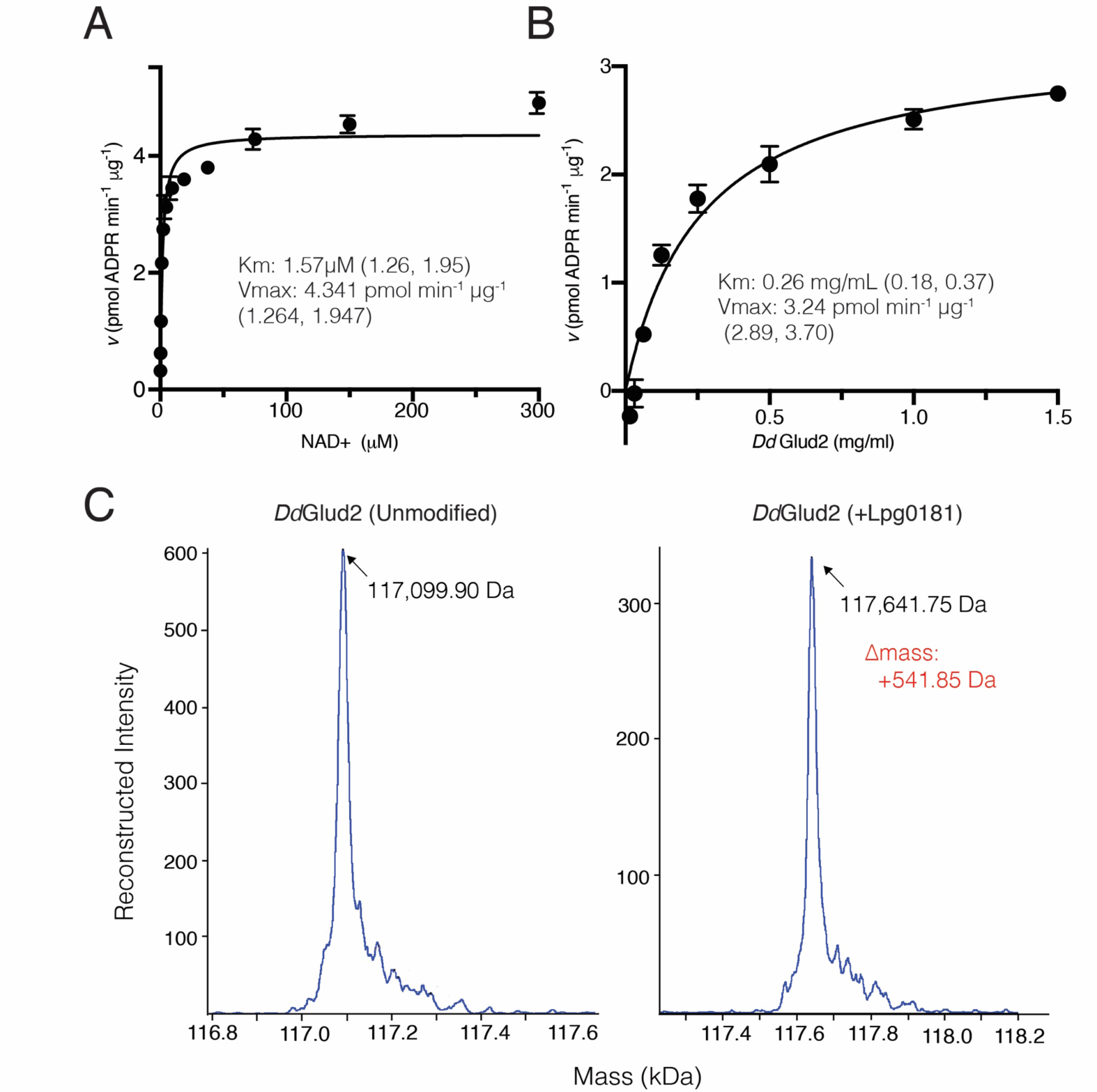
Kinetic parameters and stoichiometry of the Lpg0181 transferase reaction. **(A)** Rate plot depicting the specific activity of Lpg0181 at saturating [*Dd*Glud2] and varying [NAD+]. **(B)** Specific activity of Lpg0181 at saturating [NAD+] while varying [*Dd*Glud2]. Km and Vmax (inset) are indicated along with a 95% confidence interval. Reactions were performed in triplicate and are representative of three independent experiments. **(C)** Intact mass spectra of unmodified *Dd*Glud2 (left) or after incubation with NAD+ and Lpg0181 (right). The theoretical MW of DdGlud2 is 117,142.74 Da and the theoretical mass increase of ADP-ribosylation is 541 Da.

### Lpg0181 targets a conserved arginine residue in the nucleotide-binding pocket of GDH

To identify the modified residue of yeast Gdh2, ADP-ribosylation was enzymatically converted to phosphoribosylation by snake venom phosphodiesterase to aid identification by mass spectrometry (35). LC-MS/MS identified neutral loss of the phosphoribose group in peptides corresponding to Gdh2 791-804 (**Fig. 4*A***). Three well-conserved residues in this peptide (E792, R795, and R800) were targeted by alanine mutagenesis, and only the R800A mutant was not ADP-ribosylated by Lpg0181 (**Fig. 4*B***). Mutation of the corresponding residue (R763) in *Dd*Glud2 also abolished ADP-ribosylation by Lpg0181 (**Fig. S4**). There are presently no structures of GDHs of the 115-kDa class. However, the target Arg falls within the relatively well-conserved core nucleotide-binding domain, which can be confidently modeled based on structures of the 50 kDa-class GDHs. A model of *Dd*Glud2 shows that R763 is positioned in a solvent-exposed “lid” within the NAD+-binding pocket which closes over the ribose moiety when nucleotide is bound (**Fig. 4*C***). R763 and the surrounding motif are strongly conserved within fungal and protist GDH enzymes but have diverged in GDH classes from metazoans and bacteria (**Fig. 4*D***).

**Figure 4.**
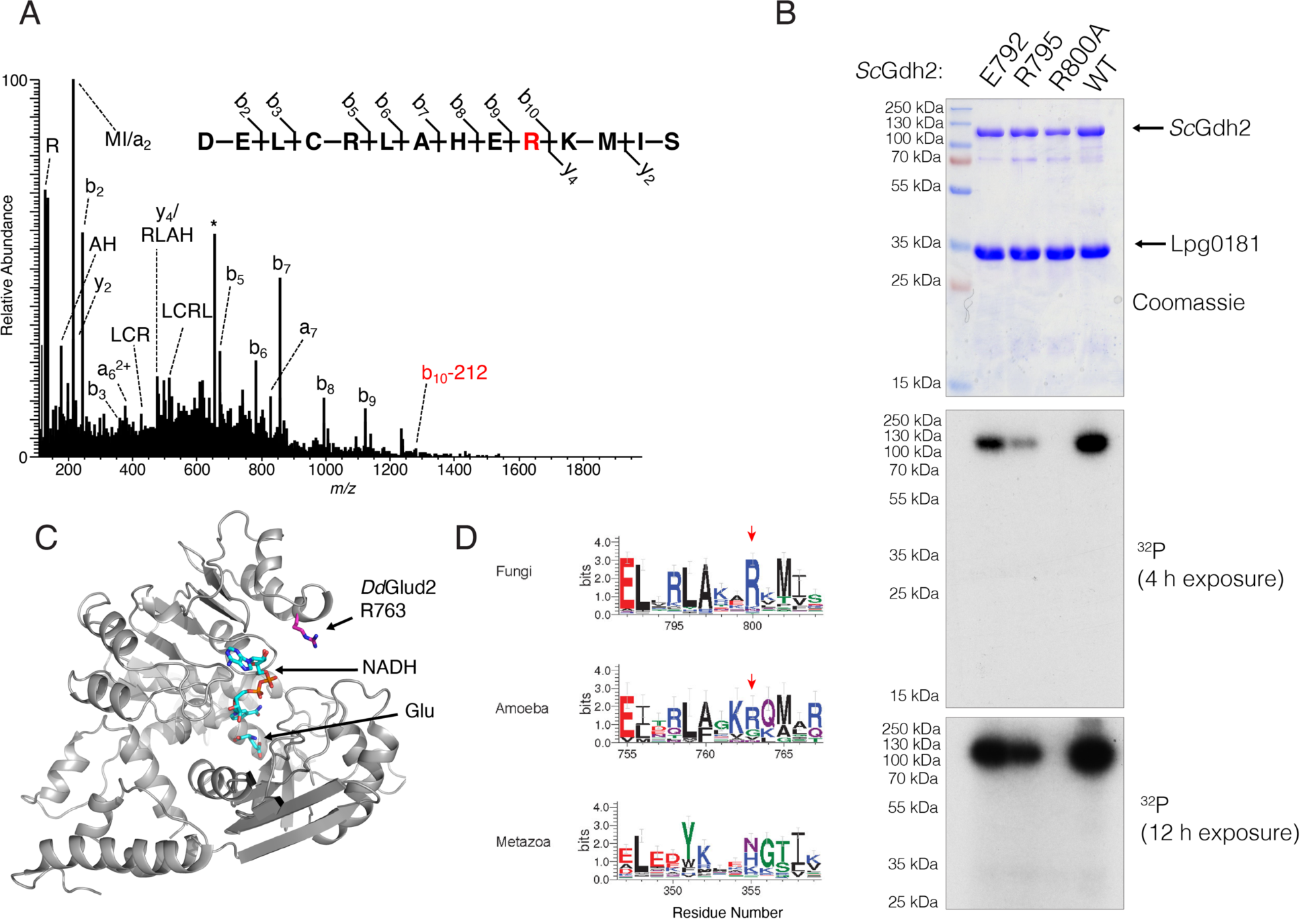
Lpg0181 targets a conserved arginine residue in the NAD+-binding pocket of glutamate dehydrogenase. **(A)** Fragmentation pattern of *Sc*Gdh2 peptide containing phosphoribosyl-Arg800 identified by LC-MS/MS. The precursor [M+3H]^3+^ ion, *m/z* 657.30, is labeled with an asterisk (*) and was subjected to HCD fragmentation to generate the spectrum shown. The modification site was localized to arginine 800 highlighted in red. The b10 fragment ion containing the modified residue shows neutral loss of the phosphoribosyl group (−212 Da). **(B)** Endpoint assays depicting incorporation of [^32^P]-NAD+ by Lpg0181 into the indicated alanine mutants of *Sc*Gdh2. Mutation of the ADPR acceptor site R800 abolishes ADP-ribosylation. **(C)** Model of *Dd*Glud2 built by Phyre (60) using *Pyrococcus furiosus* GDH (PDB 1HRD) as a template, indicating the position of the ADP-Ribose acceptor residue Arg 763 (rendered in sticks). Bound NADH and Glutamate (Glu), (modeled by alignment to liganded bovine GDH, PDB 6DHQ) are rendered in sticks. **(D)** Sequence logos depicting the conservation of R800 and the surrounding residues in GDH enzymes from Fungi (top, based on MAFFT alignment of 125 sequences), Amoeba (middle, based on MAFFT alignment of 19 sequences) and metazoans (bottom, based on MAFFT alignment of 145 sequences). Red arrows indicate the position of the arginine targeted by Lpg0181.

Yeast Gdh2 is a predominately catabolic enzyme that uses NAD+ as a cofactor and catalyzes the oxidative deamination of glutamate (30,31). Amoeba GDHs are less well-characterized and have not been purified to homogeneity, but NAD+-dependent, catabolic GDH activity has been partially purified from the cytosol of *D*. *discoidium* (36) and *A*. *castellani (*37,38). We cloned and purified the 115-kDa class GDH from the soil amoeba *D*. *discoideum*, which shares 38% sequence identity with yeast Gdh2. The recombinant enzyme has a subunit M.W. of 117.1 kDa and elutes primarily with mr of 212 kDa (**Fig. S5**), suggesting a dimer. *Dd*Glud2 reduced NAD+ in the presence of glutamic acid and oxidized NADH in the presence of ammonia and α-KG. Rate plots with varying glutamate and α-KG are shown in **Fig. S6A** and **S6B**. The rate plot for glutamate was roughly hyperbolic and indicates a Km of 18.53 mM (95% CI 10.72, 32.15). α-KG appears to elicit substrate inhibition at concentrations above 50 mM and the rate plot could not be fit to Michaelis-Menten kinetics. As expected, *Dd*Glud2 did not utilize NADP+ as a cofactor. With substrate concentrations held near Km, its rate was not affected by ATP or CTP, but was stimulated 2.5-fold by GTP (**Fig. S6*C***).

### GDH is inactivated by ADP-ribosylation

ADP-ribosylated *Dd*Glud2 (ADPR-*Dd*Glud2) was prepared in reactions with Lpg0181 and purified by gel filtration chromatography. In glutamate oxidation assays, ADPR-*Dd*Glud2 was completely inactivated; its catalytic constant was reduced to <1% that of the unmodified enzyme (**Fig. 5A**). Similarly, in reverse reactions with α-KG, ammonia and NADH, ADPR-*Dd*Glud2 lost detectable activity (**Fig. 5B**). Thus, ADP-ribosylation of host GDH by Lpg0181 potently inactivates its metabolic function, likely by occluding the NAD+-binding pocket with an ADP-ribose moiety.

**Figure 5.**
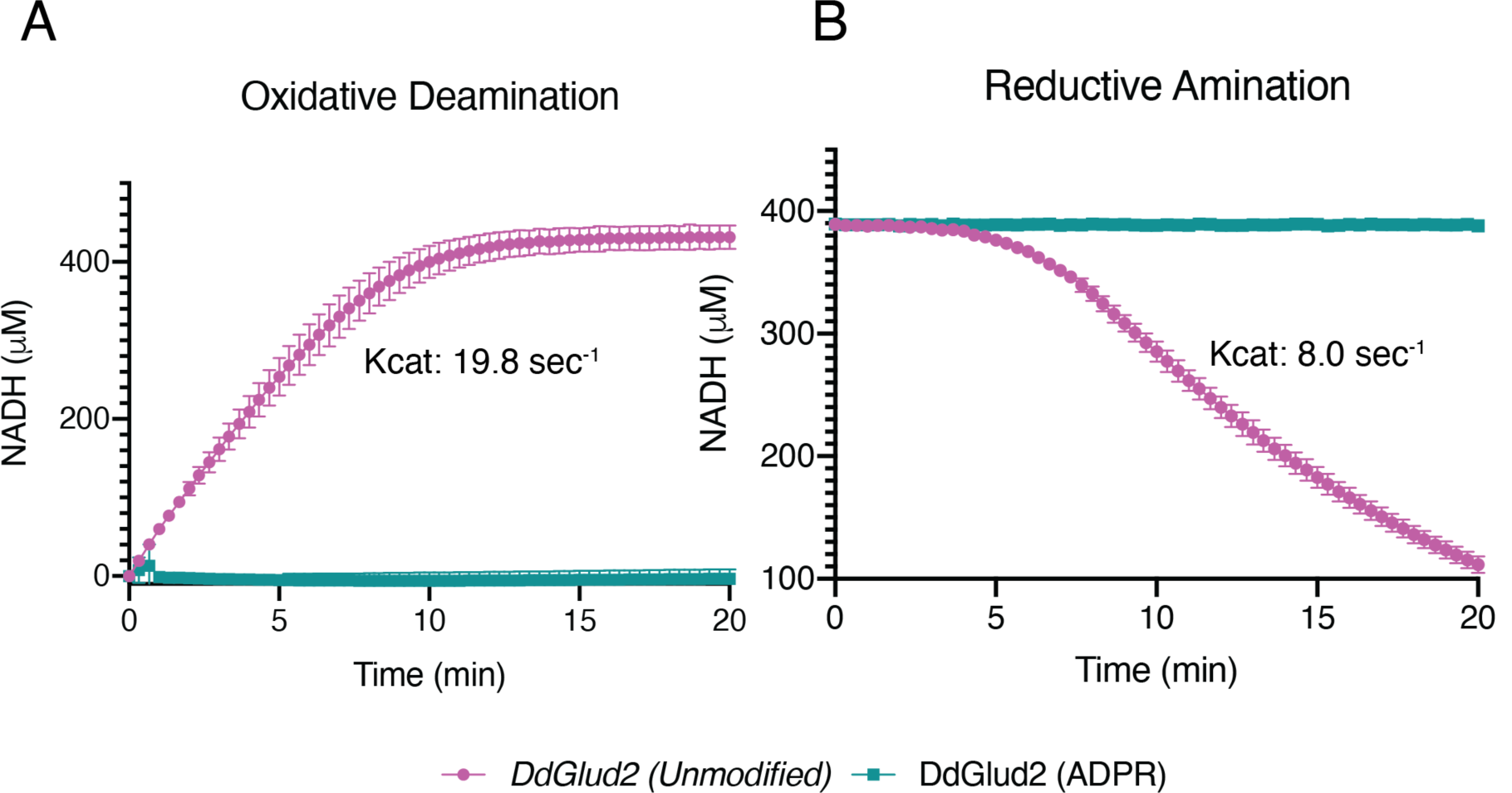
ADP-ribosylation inactivates *Dd*Glud2. Reaction progress curves showing **(A)** oxidative deamination of glutamate by unmodified (magenta) or ADP-ribosylated (teal) *Dd*Glud2, generated by detecting NAD+ reduction to NADH (Abs 340nm). Reactions contained excess NAD+ and glutamate. **(B)** Reaction progress curves showing reductive amination by unmodified (magenta) or ADP-ribosylated (teal) *Dd*Glud2. Reactions contained excess α-KG, NADH, and NH_4_^+^. Plots display mean and SD of four replicates from two independent experiments.

## Discussion

We have identified Lpg0181 as the second *Legionella* effector using ADP-ribosyltransferase activity for the pathogen’s benefit, the first being the ART domain in the SidE family of all-in-one ubiquitin ligases. The two types of ART domains are only remotely similar and perform very different functions. This highlights the pathogen’s ingenuity in evolving and repurposing specific enzymatic activities. To our knowledge, Lpg0181 is also the first *Legionella* effector identified which directly targets a host metabolic enzyme. Vertebrate GDH is subject to regulation by ADP-ribosylation, presumably as a mechanism to fine-tune insulin secretion (39-41). However, this ADP-ribosylation is catalyzed by SIRT4, which does not adopt an ART fold, occurs on a cysteine residue, and imparts reversible and partial enzyme inhibition (40,42). In contrast, Lpg0181 has evolved a distinct mechanism to completely inactivate GDH by targeting a conserved arginine residue in the NAD+-binding pocket.

We have also cloned and purified an amoeba 115-kDa NAD+-dependent GDH, extending observations of crude and partially purified NAD+-dependent GDH activity from several amoeba. While Gdh2 in yeast is thought to be a mitochondrial protein, amoeba GDH activity is present in the cytosol (36,37,43). None of our observations indicate that Lpg0181 would localize to the mitochondria, suggesting it could encounter amoeba GDH in the cytosol.

Glutamate dehydrogenases are ubiquitous enzymes that occupy a key metabolic branch point, liberating nitrogen from amino acids and supplying carbon chains to the TCA as oxoglutarate. While we were unable to determine if Lpg0181 can modulate host metabolism during infection, *Legionella* and other intracellular bacteria are entirely dependent on nutrients extracted from their host cell. The concept of ‘nutritional virulence’ postulates that intracellular pathogen virulence strategies are driven by a need to override host nutrient restrictions (44). The levels of free amino acids in host cytosol are insufficient to support the demands of replicating intravacuolar bacteria (45-47). Thus, secreted effectors probably play a key role in liberating nutrients and triggering host cell processes that increase metabolite levels. For instance, the *Legionella* effector AnkB directs widespread K48-linked ubiquitination of host proteins, which are digested by the proteasome. The resulting free amino acids accumulate and supply the dividing bacteria with macronutrients (47). Other intracellular pathogens liberate metabolites by triggering host autophagy (48). Lpg0181 may also function to increase the concentration of free glutamate by directly inactivating host glutaminolysis.

Alternatively, GDH may have additional functions targeted through Lpg0181. Amoeba glutamate dehydrogenases may have important roles in the response to osmotic stress or starvation, cues that induce encystation and restrict bacterial intracellular replication (38,43,49). Thus, Lpg0181 may target amoeba GDH to prevent a conserved stress response.

Deletion of Lpg0181 had no effect on *Legionella* replication in *A*. *castellani*, suggesting it targets a pathway which is manipulated by additional effectors **(Fig. S7)**. Because Lpg0181 specifically recognizes a subfamily of GDH enzymes only present in fungi and protists, it is unlikely to contribute to virulence in vertebrate macrophages. Lpg0181 is thus an example of an amoeba-specific ‘auxilliary’ gene that promotes *Legionella* parasitism in natural hosts and contributes to its broad host range (50).

In conclusion, our results uncover a novel member of the mono-ART family and demonstrate that *Legionella* has evolved to directly target host metabolic enzymes as part of its pathogenic strategy.

## Experimental Procedures

### Reagents

ATP (A2383), CTP, GTP, α-ketoglutaric acid (75890) L-glutamic acid (G1626), NAD+ (10127965001) NADH (10107735001) NADP+ (93205) and Protease inhibitor cocktail (PIC; 11873580001) were from Sigma. Six-biotin-17-NAD+ (4670-500-01) was from Trevigen. [^32^P]-NAD (NEG023×250UC) was from Perkin Elmer. Tris (2-carboxyethyl) phosphine hydrochloride (TCEP; 20490), PfuTurbo DNA polymerase (50-125-946) and high-capacity streptavidin agarose (20357) were purchased from Thermo Fisher Scientific. Q5 DNA polymerase, 2X Gibson assembly mix, and all restriction enzymes used for cloning were from New England Biolabs.

### Bioinformatic analysis of Lpg0181

The similarity of lpg0181 to known ADP-ribosyltransferases was established by screening the set of *Legionella pneumophila* subsp Philadelphia effectors using the FFAS server (25). Homologs of the lpg0181 ART domain and homologs of GDH isoforms were collected using BLAST searches and aligned by MAFFT (51). Sequence logos were produced using the Weblogo 3.0 server (52). The CLANS algorithm (53) was used to represent sequence similarities between nineteen ART-like families from the ADP-ribosyl clan (Pfam database identifier CL0084) and four additional novel ART-like families (lpg0181, NEURL4, SidE, EspJ). Significant and sub-significant BLAST similarities up to E-value 1 were considered. For the CLANS analysis, ART-like sequence sets were downloaded from the Pfam database reference proteomes rp15 sets (54), or (for the SidE ART domains, bacterial EspJ-like ART domains and human NEURL4-like ART domains) collected using BLAST.

### Generation of Plasmids

Lpg0181, all GDH homologs, and mutants were cloned into a modified pet28a bacterial expression vector (ppSUMO), containing an N-terminal 6X-His tag followed by the yeast SUMO (SMT3) CDS. The coding sequence of *Dd*Glud2 from the genome of *D*. *discoidium* AX4(55) was accessed from dictyBase(56). The *Dd*Glud2 gene (DDB0233691) contains a single intron and a single base (T787) which are removed from the mRNA. The two exons were amplified with NEB Q5 polymerase from *D*. *discoidium* AX2 genomic DNA and joined by gibson assembly into ppSUMO. T787 was removed by site-directed mutagenesis. The construct was confirmed by sequencing and confirmed to be identical to the CDS reported in dictyBase. *Sc*GDH1 (*DHE4*, SGD:S000005902) and *Sc*GDH2 (*DHE2*, SGD:S000002374) were amplified from BY4741 gDNA, Lpg0181, Lpg1581 and Lpg0245 (accessed from *L*. *pneumophila* genome(57) assembly GCA_000008485.1) were amplified from *Legionella pneumophila* strain Philidelpia-1 gDNA, and the *Hs*Glud2 cDNA clone (NM_012084.3) was obtained from the Ultimate™ ORF Lite human cDNA collection (Life Technologies). Amino acid mutations were introduced via Quick Change site-directed mutagenesis as previously described(58). Briefly, primers were designed using the Agilent Quick Change Primer design tool: https://www.genomics.agilent.com and used in PCR reactions to generate the desired mutation using PfuTurbo DNA polymerase. Reaction products were digested with DpnI restriction endonuclease and mutations were confirmed by sanger sequencing.

### Expression and purification of recombinant Lpg0181

ppSUMO-Lpg0181and the E137A mutant were transformed into BL21 Rosetta cells for protein expression. Cells were grown in Luria Bertani (LB) broth supplemented with kanamycin (50 *μ*g/mL) to OD_600_ of 0.8-1.1 at 37 °C with constant orbital shaking at 250 rpm. Cells were cooled to 23 °C and protein expression was induced with 0.4 mM IPTG for 16-18 hours at 23 °C with orbital shaking at 250 rpm. Cells were harvested by centrifugation at 4,000 xg for 15 min and lysed in 50 mM Tris-HCl pH 8, 300 mM NaCl, 1 mM PMSF, 1 mM DTT by sonication. Cell lysates were cleared by centrifugation at 30,000 -35,000 x g for 30 minutes. The cleared lysate was incubated with washed Ni-NTA beads for a minimum of one hour at 4°C. Beads were collected in a gravity-flow column and washed with 20 column volumes of 50 mM Tris-HCl pH 8, 300 mM NaCl, 25 mM imidazole, 1mM DTT. Proteins were eluted with 50 mM Tris-HCl pH 8, 300 mM NaCl, 300 mM imidazole, 1 mM DTT. Eluates were concentrated to ∼2.5 mL, 6X-His ULP was added, and the protein was transferred to 10,000 Da mwco cellulose dialysis tubing and dialyzed against 5L of 50 mM Tris-HCl pH 8, 300 mM NaCl, 1 mM DTT overnight at 4°C with gentle stirring. The cut protein was cleared by centrifugation at 20,000 x g for 10 minutes then the volume was increased to 20 mL with fresh dialysis buffer and incubated with fresh Ni-NTA beads for one hour at 4°C to bind the cleaved 6X-His-SUMO and 6X-His-ULP. Samples were passed over a second gravity column and the flow-through, containing Lpg0181, was collected and passed a second time over the Ni/NTA resin. The flow-through was then concentrated and further purified by gel filtration chromatography using a Superdex 75 gel filtration column in with 50 mM Tris-HCl pH 8, 300 mM NaCl, 300 mM imidazole, 1 mM DTT. Peak fractions were collected and concentrated. The purified protein was stored at 2-15 mg/mL in gel filtration buffer supplemented with 5% glycerol and flash-frozen in liquid nitrogen prior to storage at -80°C.

### Labeling of Lpg0181 substrates in yeast lysate

Yeast used in this study were BY4741[*Mata leu2Δ0 met15Δ0 ura3Δ0 his3Δ1*]. Five mL BY4741 were grown in YPD broth (20 g/L <u>peptone</u>, 10 g/L <u>yeast extract</u>, 2 %w/v glucose) for approximately 20 hours at 30 °C with orbital shaking at 250 rpm until the culture reached an OD_600_ ≥ 3. The cells were pelleted and washed twice in H_2_O by centrifugation at 800 xg for 5 minutes. The pellet was resuspended in 500 *μ*L ice-cold IP buffer (50 mM Na-HEPES, 200 mM NaOAc, 1 mM EDTA, 1 mM EGTA, 5 mM MgOAc, 5 %w/v glycerol, 0.25 %w/v NP-40, 3 mM DTT, 1 mM PMSF, Roche protease inhibitor cocktail (PIC), pH 7.5) and lysed by bead beating (vortexing, 30 s pulses x5 followed by 1 min incubations on ice) with acid-washed glass beads, followed by two subsequent spins at 3,000 xg (2 min. at 4 °C) and 20,000 xg (10 min. at 4°C). the protein concentration of the cleared lysate was measured by Bradford assay and diluted to 4 mg/mL in IP buffer.

To label substrates in yeast lysate, 20 *μ*L reactions were prepared with 40 *μ*g yeast lysate or IP buffer, 7.5 *μ*g Lpg0181 or the E137A mutant, 50 mM Tris HCl pH 6.8, and initiated by adding 100 *μ*M [^32^P]-adenylate NAD+, s.a. 3000 cpm/pmol. Reactions were incubated at 37 °C for 8 min, then stopped by addition of 5*μ*L 5X SDS-PAGE loading buffer (1x = 12.5 mM Tris-PO_4_ pH 6.8, 10% (w/v) glycerol, 1.25% (w/v) SDS, 0.02% (w/v) bromophenol blue) with β-ME (1% final) and boiled for 10 min. Reactions were resolved by SDS-PAGE and ^32^P incorporation was detected by autoradiography.

### Streptavidin capture of Lpg0181 substrate

Biotin labeling was performed in 100 *μ*L reactions containing 0.3 mg yeast lysate, 0.8 *μ*g Lpg0181 or the E137A mutant, and 50 *μ*M 6-biotin-17-NAD+ (Trevigen). Reactions were incubated at 37 °C for 15 minutes, then the volume was increased to 1 mL with ice-cold 1 mM NAD+ in 50 mM Tris HCl pH 7.5. Biotin-labeled proteins were enriched following an adapted BioID protocol(59). Streptavidin agarose resin (Pierce) was washed in 50 mM Tris HCl pH 7.5 and 20 *μ*L resin volume was added to the samples. Samples were nutated at 4 °C for 1 h. The resin was collected by centrifugation (1000 xg x 2 min) and washed twice in 1 mL 2 %w/v SDS, then once in buffer 2 (0.1 %w/v deoxycholic acid, 1 %w/v Triton X-100, 1 mM EDTA, 50 mM HEPES pH 7.5) and once in buffer 3 (0.5 %w/v deoxycholic acid, 0.5 %w/v NP-40, 1 mM EDTA, 250 mM LiCl, 10 mM Tris HCl pH 7.4). The beads were then collected and suspended in a final volume of 100 *μ*L Tris HCl pH 7.5 and submitted for protein identification by LC-MS/MS.

### Protein identification by LC-MS/MS

An aliquot of streptavidin beads was incubated with trypsin at 37°C overnight to elute bound proteins. Resulting tryptic peptides were de-salted via solid phase extraction (SPE) prior to mass spectrometry analysis. LC-MS/MS experiments were performed on a Thermo Scientific EASY-nLC liquid chromatography system coupled to a Thermo Scientific Orbitrap Fusion Lumos mass spectrometer. Peptides were loaded onto a C18 column (75 *μ*m ID x 50 cm, 2 *μ*m particle size, 100Å pore size) (Thermo Fisher Scientific) and eluted with a gradient: 0-5% B in 5 min, 5-30% B in 65 min, 30-60% B in 10 min, 60-100% B in 8 min. Buffer A consisted of 2% (v/v) acetonitrile and 0.1% formic acid in water. Buffer B consisted of 80% (v/v) acetonitrile, 10% (v/v) trifluoroethanol, and 0.1% formic acid in water. To generate MS/MS spectra, MS1 spectra were first acquired in the Orbitrap mass analyzer (resolution 120,000). Peptide precursor ions were then isolated and fragmented using high-energy collision-induced dissociation (HCD). The resulting MS/MS fragmentation spectra were acquired in the ion trap. MS/MS spectral data was searched using Mascot 2.5 (Matrix Science). Precursor and fragment ion tolerances of 15 ppm and 0.6 Da, respectively, were specified and three missed cleavages were allowed. Oxidation of methionine (+15.995 Da) and carbamidomethylation of cysteine residues (+57.021 Da) were set as variable modifications.

### Purification of *Dd*Glud2

ppSUMO-*Dd*Glud2 was transformed into BL21 Rosetta cells for protein expression. Cells were grown in Luria Bertani (LB) broth supplemented with kanamycin (50 *μ*g/mL) to OD_600_ of 0.8-1.1 at 37 °C with constant orbital shaking at 250 rpm. Cells were cooled to 18 °C and protein expression was induced with 0.4 mM IPTG for 16-18 hours at 18 °C with orbital shaking at 250 rpm. Cells were harvested by centrifugation at 4,000 xg for 15 min and lysed in 50 mM Tris-HCl pH 7.5, 150 mM NaCl, 1 mM PMSF, 1 mM DTT by sonication. Cell lysates were cleared by centrifugation at 30,000 -35,000 x g for 30 minutes. The cleared lysate was incubated with washed Ni-NTA beads for a minimum of one hour at 4°C. Beads were passed over a column and washed with 20 column volumes of 50 mM Tris-HCl pH 7.5, 150 mM NaCl, 25 mM imidazole, 1mM DTT. Proteins were eluted with 50 mM Tris-HCl pH 7.5, 150 mM NaCl, 300 mM imidazole, 1 mM DTT. Proteins were cut overnight at 4°C with 6X-His tagged ULP Sumo protease followed by gel filtration chromatography using a Superdex 200 gel filtration column attached to an AKTA Pure FPLC chromatography system (GE Healthcare). Peak fractions were collected and concentrated. The purified protein was stored at 2-20 mg/mL in gel filtration buffer protected with 5% glycerol and flash-frozen in liquid nitrogen prior to storage at -80°C.

### Purification of other GDH homologs

*Hs*Glud2, Lpg1581, Lpg0245 *Sc*GDH1, *Sc*GDH2 and mutants of *Sc*GDH2 were purified as described for *Dd*GLud2, except that protein expression was induced at 23 °C, and buffers contained 50 mM Tris-HCl pH 8, 300 mM NaCl, 1 mM PMSF, 1 mM DTT.

### Preparation of ADPR-*Dd*Glud2

ADP-ribosylation was performed in a 100 *μ*L reaction containing 0.5 mg *Dd*Glud2, 0.1 mg Lpg0181, 10 mM NAD+, 50 mM Tris HCl pH 8.0, 150 mM NaCl, 1mM TCEP for 1.5 h at 23°C. Following a 10 min spin at 10,000 xg to remove aggregated protein, the reaction was cooled to 4°C and separated on a Superdex 200 increase gel filtration column equilibrated in 50 mM Tris HCl pH 8.0, 150 mM NaCl, 1mM TCEP. The fractions containing *Dd*Glud2 were collected, concentrated, and submitted for intact mass analysis or used in activity assays.

### Intact mass analysis of ADPR-*Dd*Glud2 (AL)

Unmodified and ADP-ribosylated *Dd*Glud2 prepared as described above were analyzed using a Sciex X500B Q-ToF mass spectrometer coupled to an Agilent 1290 Infinity II HPLC. Samples were injected onto a POROS R1 reverse-phase column (2.1 x 30 mm, 20 *μ*m particle size, 4000 Å pore size), desalted, and the amount of buffer B was manually increased stepwise until the protein eluted off the column. Buffer A contained 0.1% formic acid in water and buffer B contained 0.1% formic acid in acetonitrile. The mobile phase flow rate was 300 μL/min.

The mass spectrometer was controlled by Sciex OS v.1.3 using the following settings: Ion source gas 1 15 psi, ion source gas 2 30 psi, curtain gas 35, CAD gas 7, temperature 200 °C, spray voltage 5200 V, declustering potential 80 V, collision energy 15 V. Data was acquired from 1400-3600 Da with a 1 s accumulation time and 80 time bins summed. The acquired mass spectra for the proteins of interest were deconvoluted using BioPharmaView v. 2.1 software (Sciex) in order to obtain the molecular weights. The peak threshold was set to ≥ 5%, reconstruction processing was set to 20 iterations with a signal to noise threshold of ≥ 5 and a resolution of 20000.

### LC-MS/MS Analysis of phospho-ribosylated peptides in *S*. *cerevisiae* Gdh2

Gdh2 was ADP-ribosylated in 20 *μ*L reactions with 0.5 mg/mL Gdh2, 0.125 Lpg0181, 50 mM Tris HCl pH 7.5, 150 mM NaCl, 1 mM DTT, and 2.5 mM NAD+. After a 20 min incubation at 37 °C, 1.5 *μ*L of 1M Tris HCl pH 9.0, 0.5 *μ*L of 0.5M MgCl_2_, and 5 *μ*L of 1 mg/mL snake venom phosphodiesterase I (Sigma) was added to convert ADP-ribosylation to phospho-ribosylation. The reaction was incubated an additional hour at 37 °C then the reaction was boiled in 1X SDS-PAGE + β-ME sample buffer. The entire sample was resolved by SDS-PAGE. The band corresponding to Gdh2 was excised with a razor and submitted for mass spectrometry.

Samples were reduced with DTT and alkylated with iodoacetamide prior to overnight enzymatic digestion with Asp-N at 37°C. Resulting peptides were de-salted via solid phase extraction (SPE) prior to LC-MS/MS analysis. Experiments were performed on a Thermo Scientific EASY-nLC liquid chromatography system coupled to a Thermo Scientific Orbitrap Fusion Lumos mass spectrometer in the same way described above. Samples were searched using Mascot 2.5 (Matrix Science). Precursor and fragment ion tolerances of 15 ppm and 0.6 Da, respectively, were specified and three missed cleavages were allowed. Oxidation (M) (+15.995 Da), carbamidomethylation (C) (+57.021 Da), and phospho-ribosylation (DER) (212.009 Da) were set as a variable modifications. MS/MS spectra of phospho-ribosylated peptides identified by Mascot were verified manually.

### ADP-ribosylation assays

Reactions were typically conducted in 20 *μ*L volumes with 50 mM Tris HCl pH 6.8, 50 mM NaCl, 1mM DTT, contained 0.2 – 0.5 mg/mL GDH substrates, and were initiated with 100 *μ*M [^32^P]-adenylate NAD+, s.a. 500 -1000 cpm/pmol. Lpg0181 was added to 0.005 mg/mL for the timecourse, 0.03 mg/mL for the substrate panel, and 0.25 mg/mL for the alanine mutants of *Sc*Gdh2. Reactions were conducted at 37 °C for 15 - 20 min or as indicated, then quenched with 2 *μ*L 50 mM unlabeled NAD+ (pH 8) then 5X SDS-PAGE sample buffer + β-ME was added and the samples were boiled for 10 min. Products were resolved by SDS-PAGE and stained with Coomassie, dried in a gel dryer, and ^32^P incorporation was detected by autoradiography.

To determine the salt and pH optimum for Lpg0181, 20 *μ*L reactions were performed in 100 mM Sodium acetate, 50 mM Bis-Tris, 50 mM Tris-HCl (pH series) and NaCl from 0 to 500 mM. Reactions contained 0.5 mg/mL *Dd*Glud2, 0.007 mg/mL Lpg0181, and were initiated with 100 *μ*M [^32^P]-adenylate NAD+, s.a. 750 cpm/pmol. Reactions were conducted at 23 °C for 20 min. To determine the Km for NAD+, 20 *μ*L reactions contained 0.5 mg/mL *Dd*Glud2 with 0.007 mg/mL Lpg0181 in 50 mM Tris HCl pH 7.5, 150 mM NaCl, 1 mM DTT. Reactions (in triplicate) were initiated with [^32^P]-adenylate NAD+, s.a. 750 cpm/pmol at thirteen dilutions from 0 to 300 *μ*M and allowed to proceed at 23 °C for 20 min. To determine the Km for *Dd*Glud2, reactions were performed as above except Lpg0181 was used at 0.003 mg/mL and [^32^P]-NAD+ was held at 100 *μ*M (s.a. 1000 cpm/pmol) while *Dd*Glud2 was varied from 0.0156 to 1.5 mg/mL. Reactions were stopped with 5 *μ*L of a 5X stop mix containing 80 mM NAD+ in 5X SDS-PAGE sample buffer + β-ME, pH 6.8, boiled 10 min, and resolved by SDS-PAGE. Background was determined in samples that were boiled in stop mix before the addition of [^32^P]-NAD+ mix. Gels were stained with Coomassie blue, washed extensively with water and destain solution to remove background signal, then the *Dd*Glud2 bands were excised with a razor and transferred to scintillation vials. Background radioactivity (typically ∼50 cpm, less than 10% of the lowest experimental value) was subtracted from each measurement. Rate measurements were fit to Michaelis-Menten kinetic models and Km and V_max_ for substrates were calculated by nonlinear regression using Prism 8.4.1 for macOS (GraphPad Software, San Diego, California USA, www.graphpad.com).

### ADP-ribosylation assays with ethenoadenosine-NAD+

Reactions were conducted in 25 *μ*L volumes with 50 mM Bis-Tris HCl pH 6.5, 50 mM NaCl, 1mM DTT, contained 0.2 mg/mL *Dd*Glud2 substrate, 250 *μ*M 1, N^6^-etheno-NAD+, and were initiated with 0.625 *μ*g Lpg0181 or the indicated ART enzyme. Some reactions contained 500 *μ*M ADP-ribose where indicated. After a 30 min incubation at 37°C, reactions were quenched by addition of unlabeled NAD+ to 5 mM and terminated by boiling in 1X SDS-PAGE loading buffer with 1% β-ME. Immunoblotting for ethenoadenosine was performed with 1G4 (sc-) diluted in 2% nonfat milk in TBS-T.

### Modeling *Dd*Glud2

The entire *Dd*Glud2 protein sequence was submitted for modeling by Phyre(60). 422 residues (40% of the sequence) were modelled with 100% confidence using the highest scoring template (Pyrococcus furiosus glutamate dehydrogenase, PDB 1HRD, 22% identity). The binding sites for Glu, NAD+ were determined by superposition of the model with ligand-bound bovine GDH (PDB 6DHQ).

### Glutamate oxidation and ammonia assimilation assays for *Dd*Glud2

Glutamate dehydrogenase activity was measured by continuously monitoring the reduction of NAD+ to NADH spectrophotometrically at 340 nm. Reactions (1 mL volume) contained 5 *μ*g *Dd*Glud2, 100 mM Tris-HCl pH 8.0, 0.1 mg/mL BSA, and 1mM DTT. NAD+ was held at 5 mM to determine the Km for Glu. NAD+ stock solutions were prepared by dissolving NAD+ (free acid) in H_2_O and adjusting the pH to between 7 and 8 with NaOH, then the concentration was calculated by measuring A_259_ with a molar extinction coefficient of 16,900 l x M^-1^ x cm^-1^ in a quartz cuvette. Glutamic acid was dissolved directly in H_2_O. Reactions were prepared at 9/10 volume without Glu, then aliquoted into clear plastic cuvettes (path length 1 cm). The reaction was initiated by adding Glu (1/10 volume) and pipetting vigorously before initiating measurements. All reactions were performed at 23 °C. Each run was blanked to a cuvette containing reaction mix without enzyme. Under these conditions the reaction rate was linear between 2 and 5 min and within the sensitivity range of the instrument (A_340_ < 2). [NADH] was determined from A_340_ using the molar extinction coefficient of NADH at 340 nm (6,300 l x M^-1^ x cm^-1^). Rate measurements were fit to Michaelis-Menten kinetic models and Km and V_max_ for substrates were calculated by nonlinear regression using Prism 8.4.1 for macOS (GraphPad Software, San Diego, California USA, www.graphpad.com). ADPR-*Dd*Glud2 was compared to unmodified *Dd*Glud2 in reactions with 5 *μ*g/mL enzyme and saturating substrate concentrations. In some reactions, NADP+ was substituted for NAD+, or ATP, GTP, and CTP were added at 1 mM final with 10 mM Glu and 1 mM NAD+.

To measure ammonium assimilation, α-ketoglutarate (α-KG) stock was prepared by dissolving α-ketoglutaric acid (Sigma) in H_2_O and fresh NH_4_^+^ stock was prepared by neutralizing NH_4_OH to pH 8 with HCl. Reactions contained 5 *μ*g *Dd*Glud2, 100 mM Tris HCl pH 8.0, 0.1 mg/mL BSA, and 1mM DTT. NH_4_^+^ was held at 100 mM and NADH was held at 1 mM while α-KG was varied. The linear portion of each reaction was used to calculate rate plots. ADPR-*Dd*Glud2 was compared to unmodified *Dd*Glud2 in reactions with 100 mM NH_4_^+^ and 40 mM α-KG.

### Generation of ΔLpg0181 *Legionella* strain

*L*. *pneumophila* strains Lp02, Lp03 (Lp02 Δ*dotA*), and thymidine auxotrophic derivatives used in this study were derived from *Legionella pneumophila* Philadelphia-1 strain(61) and were generous gifts from Dr. Ralph Isberg. *Legionella* bacteria were maintained on ACES [*N*-(2-acetamido)-2-aminoethanesulfonic acid]-buffered charcoal yeast extract (CYE) agar plates or grown in ACES-buffered yeast extract (AYE) liquid cultures supplemented with ferric nitrate (0.135 g/L) and cysteine (0.4 g/L). Thymidine was added to a final concentration of 100 μg/mL for maintenance of the thymidine auxotrophic strains. Lpg0181 knockout strains were generated using the R6K suicide vector pSR47s (Kan^R^, *sacB*)(62), a generous gift from Dr. Shaeri Mukherjee, UCSF. Briefly, ∼800bp regions flanking the Lpg0181 ORF (Fig. S7) were amplified and cloned using Gibson assembly into pET-21a(+), then subcloned into pSR47s to generate pSR47s-*ΔLpg0181*, which was maintained in S17-1 γpir *E*. *coli*. pSR47s-*ΔLpg0181* was introduced by electroporation into strain Lp02 and colonies having undergone homologous recombination were selected with kanamycin (20 μg/mL). Metrodiploids were resolved on 10% sucrose, and the resulting colonies were screened for loss of Lpg0181 by PCR and protein immunoblotting. Lpg0181 complementing strains were generated using the RSF1010 cloning vector pJB908 (Amp^R^ *td*Δ*i*)(63), a generous gift from Dr. Ralph Isberg. Transformants were selected on CYE medium without thymidine and complementation was verified by PCR and protein immunoblotting.

### Production of Lpg0181 antibodies

Untagged WT Lpg0181 was purified as described above and used to inoculate rabbits for generation of rabbit anti-serum (Cocalico Biologicals). Total IgG was partially purified by ammonium sulfate precipitation (64) and the α-Lpg0181 antibody was affinity-purified by coupling recombinant Lpg0181 to a HiTrap NHS-activated HP column essentially as described (65). Antibodies were concentrated, aliquoted and stored at -20°C until use at a 1:2,000 dilution in 2% nonfat milk-TBST.

### Intracellular replication in amoeba

*Acanthamoeba castellanii* was maintained as a monolayer culture in PYG medium (20 g/L protease peptone, 1 g/L yeast extract, 150 mM glucose, 4 mM Mg_2_SO_4_, 0.4 mM CaCl_2_, 0.1% (w/v) sodium citrate dihydrate, 0.05 mM Fe(NH_4_)_2_ (SO_4_)_2_ x 6H_2_O, 2.5 mM NaH_2_PO_3_, 2.5 mM K_2_HPO_3_ pH 6.5) in tissue culture flasks at 23°C. 18 hours prior to infection, confluent amoeba monolayers were collected by pipetting in ice-cold PBS, resuspended in fresh PYG, counted, and 6×10^5^ cells were seeded into individual wells of 24-well plates. 1h prior to infection, amoeba were carefully washed twice, the medium was replaced with *A*. *castellanii* buffer(66) (4 mM magnesium sulfate, 0.4 mM CaCl_2_, 0.1% (w/v) sodium citrate dihydrate, 0.05 mM Fe(NH_4_)_2_ (SO_4_)_2_ x 6H_2_O, 2.5 mM NaH_2_PO_3_, 2.5 mM K_2_HPO_3_ pH 6.5) and the plates were equilibrated at 37°C. All subsequent incubations were performed at 37°C. *Legionella* cultures at post-exponential phase were diluted in *A*. *castellanii* buffer and ∼6×10^4^ bacteria were added to each well for a multiplicity of infection (MOI) of 0.1. Infections were synchronized by centrifugation at 880 x g for 5 minutes. Infections were allowed to proceed for 1 hour, then extracellular bacteria were removed by washing each well 3 times in *A*. *castellanii* buffer before adding *A*. *castellanii* buffer to a final volume of 0.5 mL/well. At timepoints 1h, 24h, and 48h, infected Amoeba cells were lysed in 0.05% saponin in H_2_O. Serial dilutions of the infectious inoculum and the amoeba lysate were plated on CYE plates to confirm the MOI and assess bacterial growth.

## Acknowledgements

We thank members of the Tagliabracci lab for valuable input and Dr. Andrew Lemoff (UTSW Proteomics Core Facility) for help with intact mass spectrometry. This work was supported by National Institutes of Health Grants R00DK099254 and DP2GM137419 (to V. S. T.), F30HL143859-01 (to M.H.B.), Welch Foundation Grants I-1911 (to V. S. T.), and the Polish National Agency for Scientific Exchange Scholarship PPN/BEK/2018/1/00431 (to K. P.). V.S.T. is the Michael L. Rosenberg Scholar in Medical Research, a Cancer Prevention Research Institute of Texas Scholar (RR150033) and a Searle Scholar.

## Conflict of Interest

The authors declare that they have no conflicts of interest with the contents of this article. The content is solely the responsibility of the authors and does not necessarily represent the official views of the National Institutes of Health.

## Author contributions

M.H.B. and V.S.T. designed the experiments. M.H.B., A.O. and V.S.T. conducted the experiments. M.G. and K.P. performed the bioinformatics. K.A.S performed the mass spectrometry. M.H.B. and V.S.T wrote the manuscript with input from all authors.

## Supporting Information

**Figure S1.**
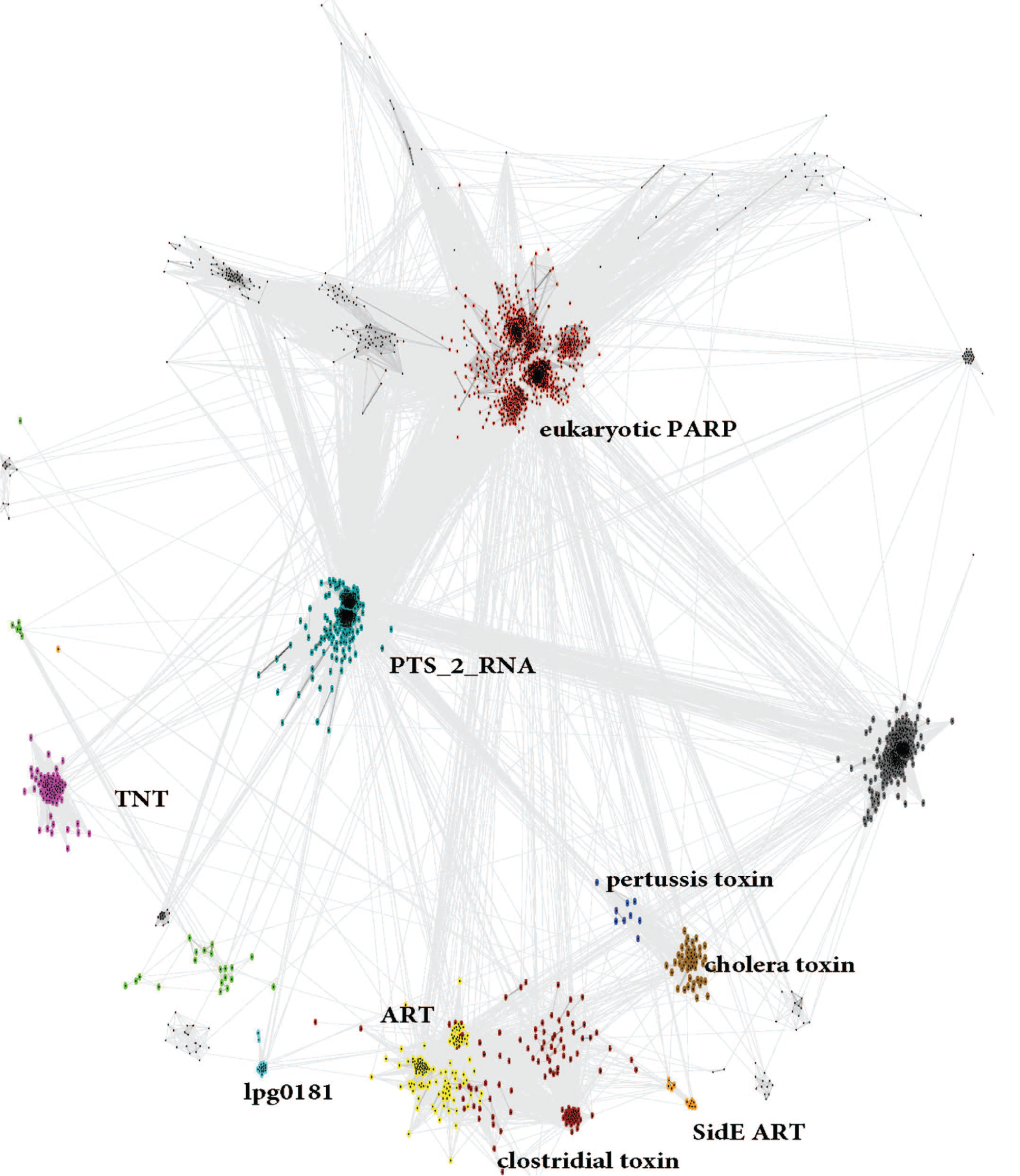
CLANS graph showing sequence similarity-based clustering of ART domain families. Light blue: lpg0181 homologs, red: PARP, yellow: ART, brown: heat-labile enterotoxin a, dark blue: pertussis toxin, cyan: PTS_2_RNA, orange: ART domains of SidE.

**Figure S2.**
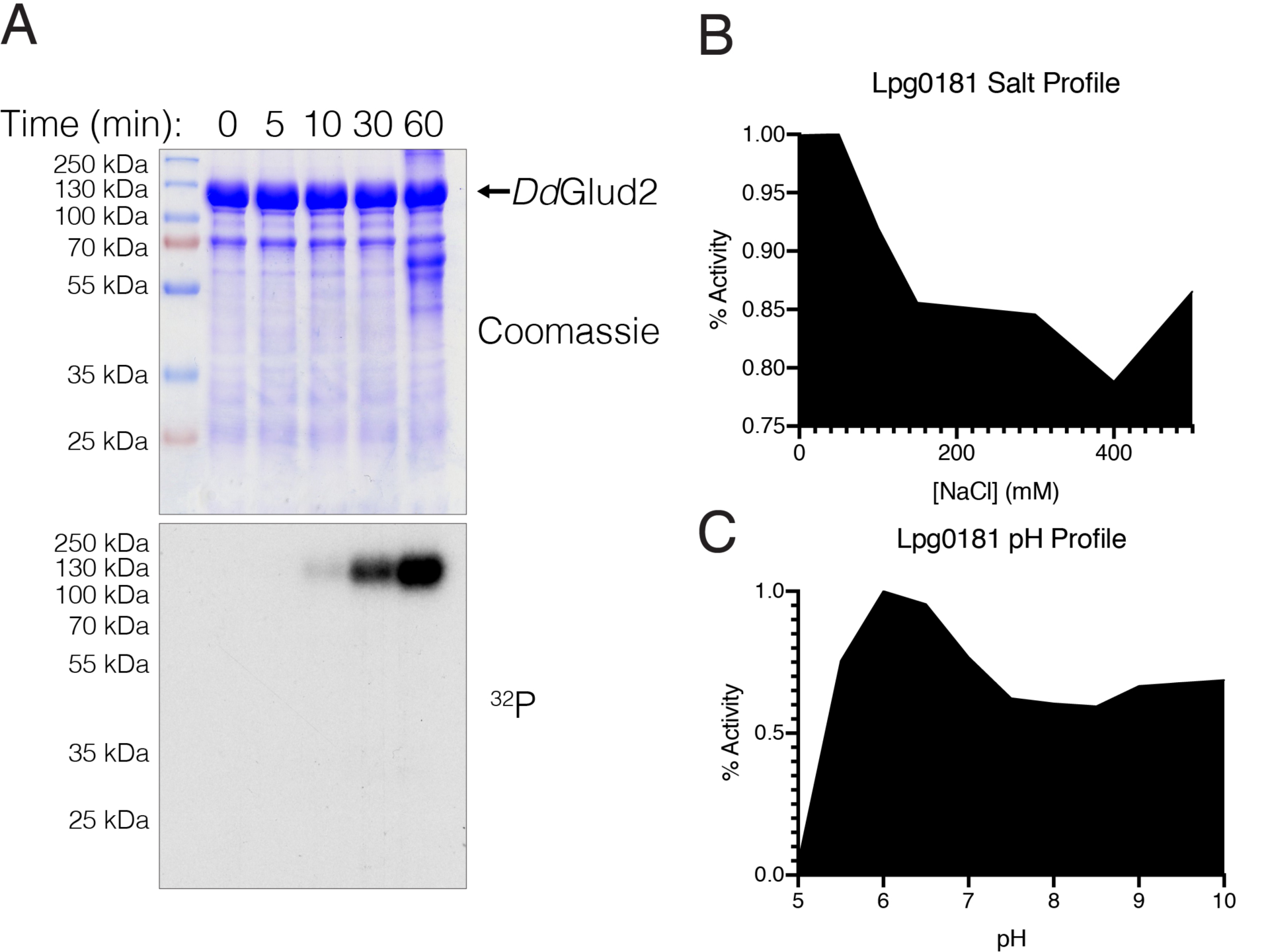
Characteristics of Lpg0181 ADP-Ribosylation. **(A)** Timecourse *in vitro* ART assay showing incorporation of ^32^P from [^32^P]-adenylate NAD+ into *Dd*Glud2 by Lpg0181. products were resolved by SDS-PAGE (upper) and visualized by autoradiography (lower). **(B)** incorporation of [^32^P]-ADPR into *Dd*Glud2 by Lpg0181 with varying concentrations of NaCl or **(C)**, buffered at the indicated pH. Reactions were resolved by SDS-PAGE and stained with Coomassie, then radioactive gel bands were excised for scintillation counting.

**Figure S3.**
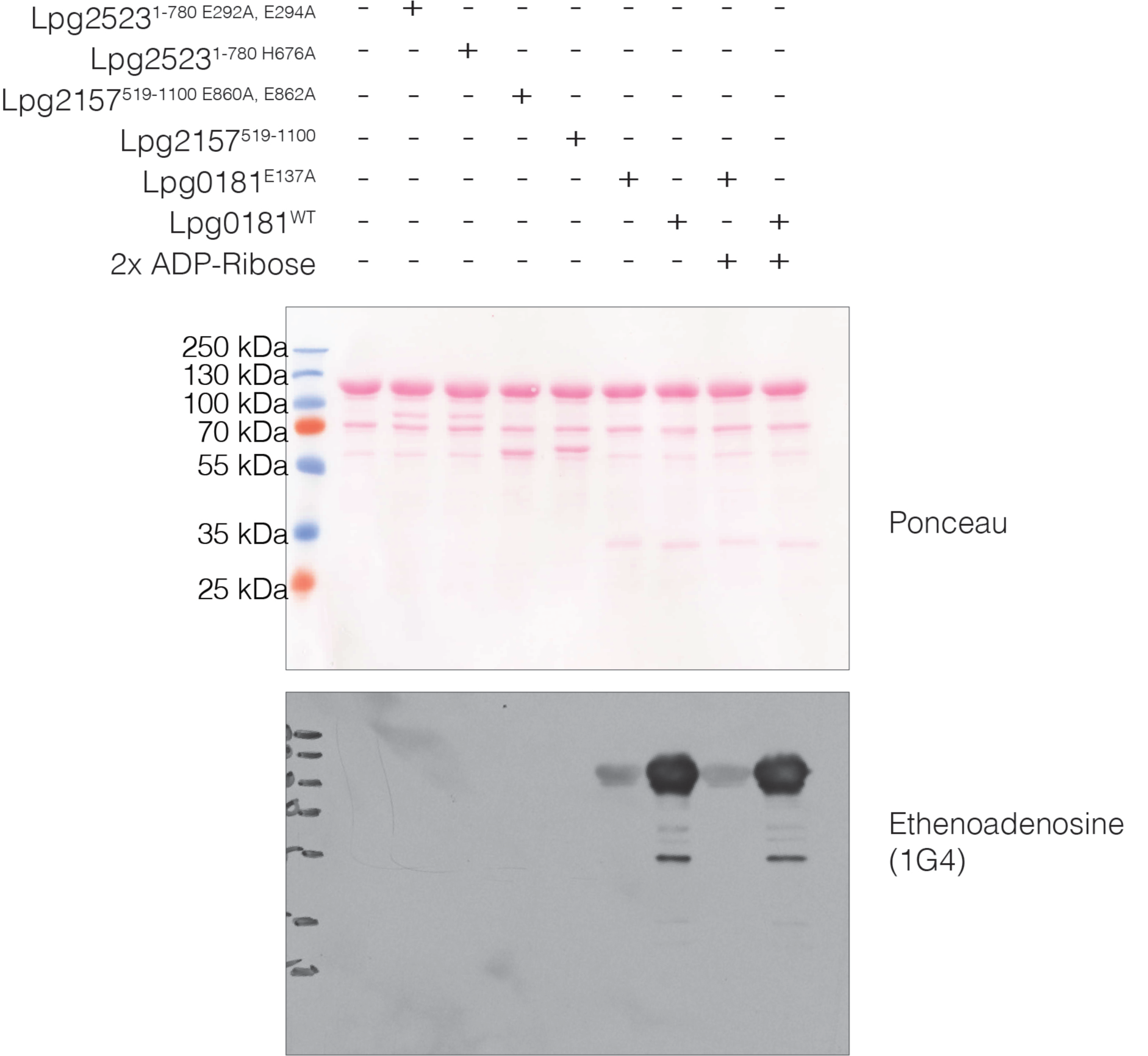
ADP-ribosylation of *Dd*Glud2 is enzyme-catalyzed. ADP-ribosylation reactions with *Dd*Glud2 as substrate were performed with nicotinamide 1,N6-ethenoadenine dinucleotide (etheno-NAD+) with no ART enzyme (lane 1), the Lpg2526 ART domain-containing effector Lpg2523 H676A (lane 2) or its ART catalytic mutant (lane 4), the SdeA (Lpg2157) ART domain and its catalytic mutant (lanes 5 and 6), or Lpg0181 and the catalytic mutant (lanes 7 and 8). In lanes 9 and 10, unlabeled ADP-ribose was added at two-fold excess to etheno-NAD+ as a test for non-enzymatic glycation. Reaction products were separated by SDS-PAGE, transferred to nitrocellulose and visualized by Ponceau staining, then etheno-ADP-ribose incorporation was detected by immunoblotting with the ethenoadenosine antibody 1G4 (Santa Cruz).

**Figure S4.**
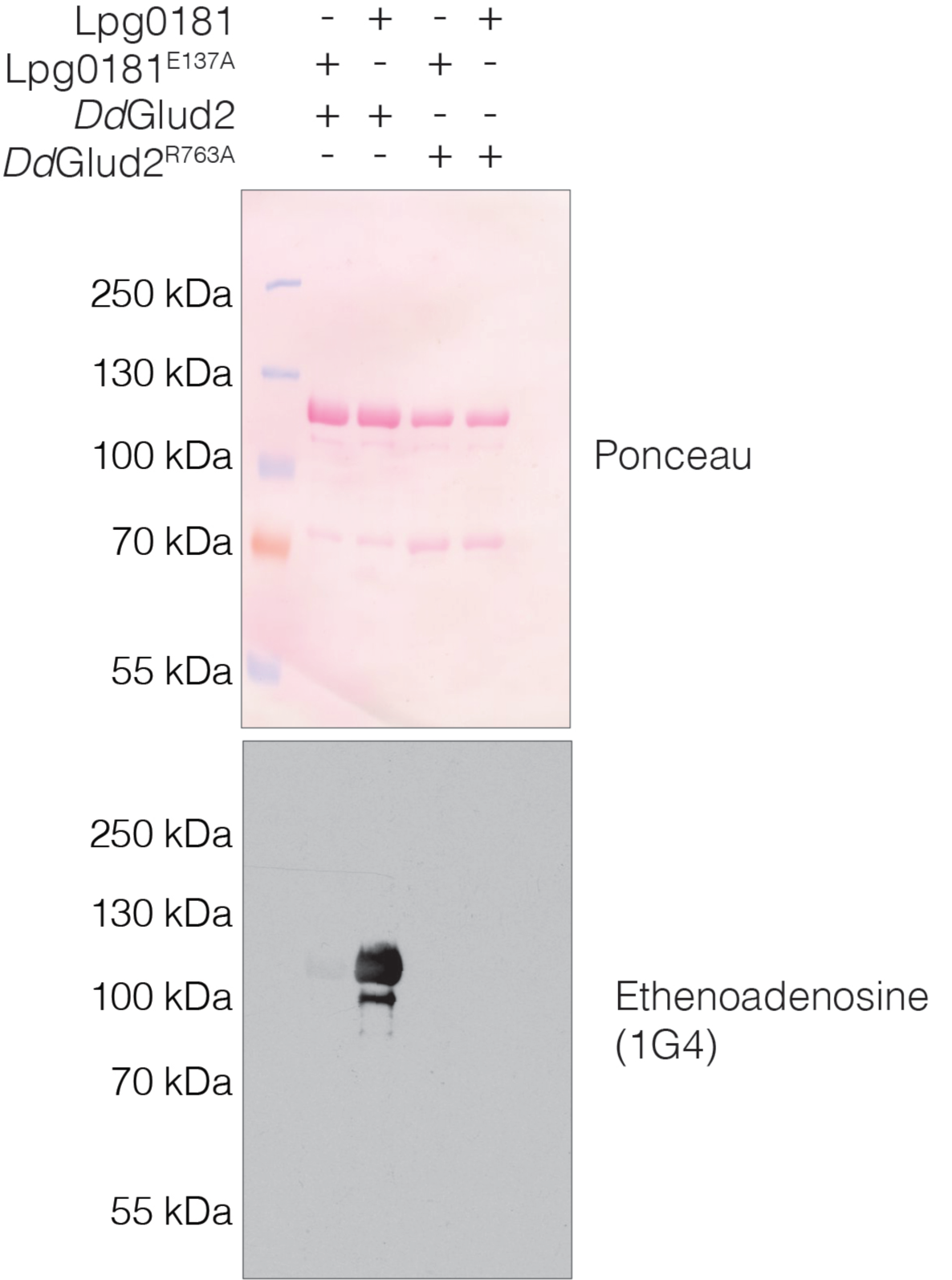
Lpg0181 targets R763 of *Dd*Glud2. ADP-ribosylation reactions with *Dd*Glud2 or *Dd*Glud2^R763A^ as substrates were performed with etheno-NAD+ and Lpg0181 or the E137A mutant. Reaction products were separated by SDS-PAGE, transferred to nitrocellulose and visualized by Ponceau staining, then etheno-ADP-ribose incorporation was detected by immunoblotting with the ethenoadenosine antibody 1G4 (Santa Cruz).

**Figure S5.**
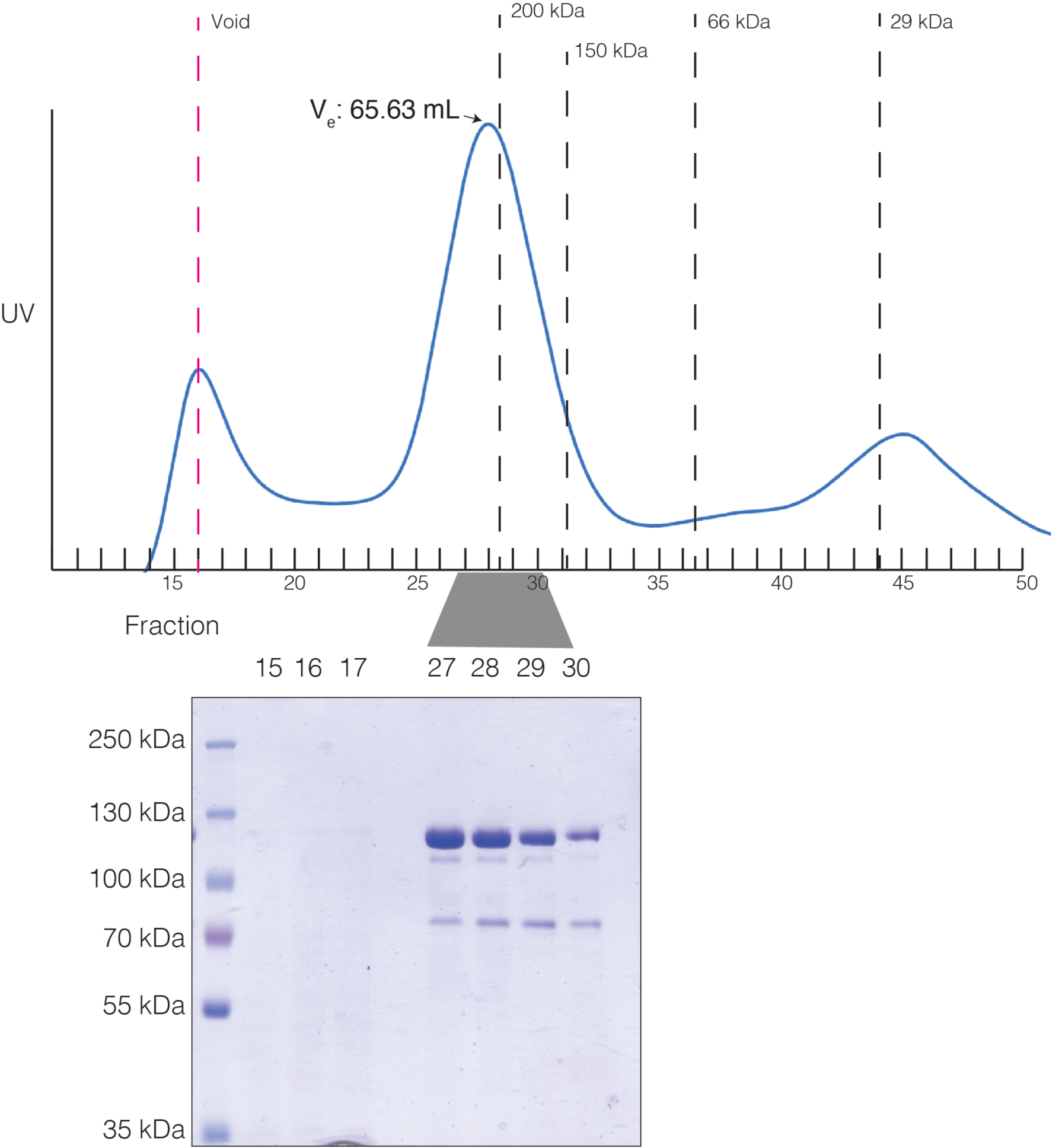
Size-exclusion chromatography of *Dd*Glud2. Recombinant *Dd*Glud2 was subjected to chromatography on a Superdex 200 column (GE). The UV trace (above) is superimposed on peak elution volumes of molecular weight standards (dotted black lines) and void elution volume (blue dextran, red dotted line). Fractions corresponding the void and peak elution were separated by SDS-PAGE and visualized by Coomassie staining (lower panel).

**Figure S6.**
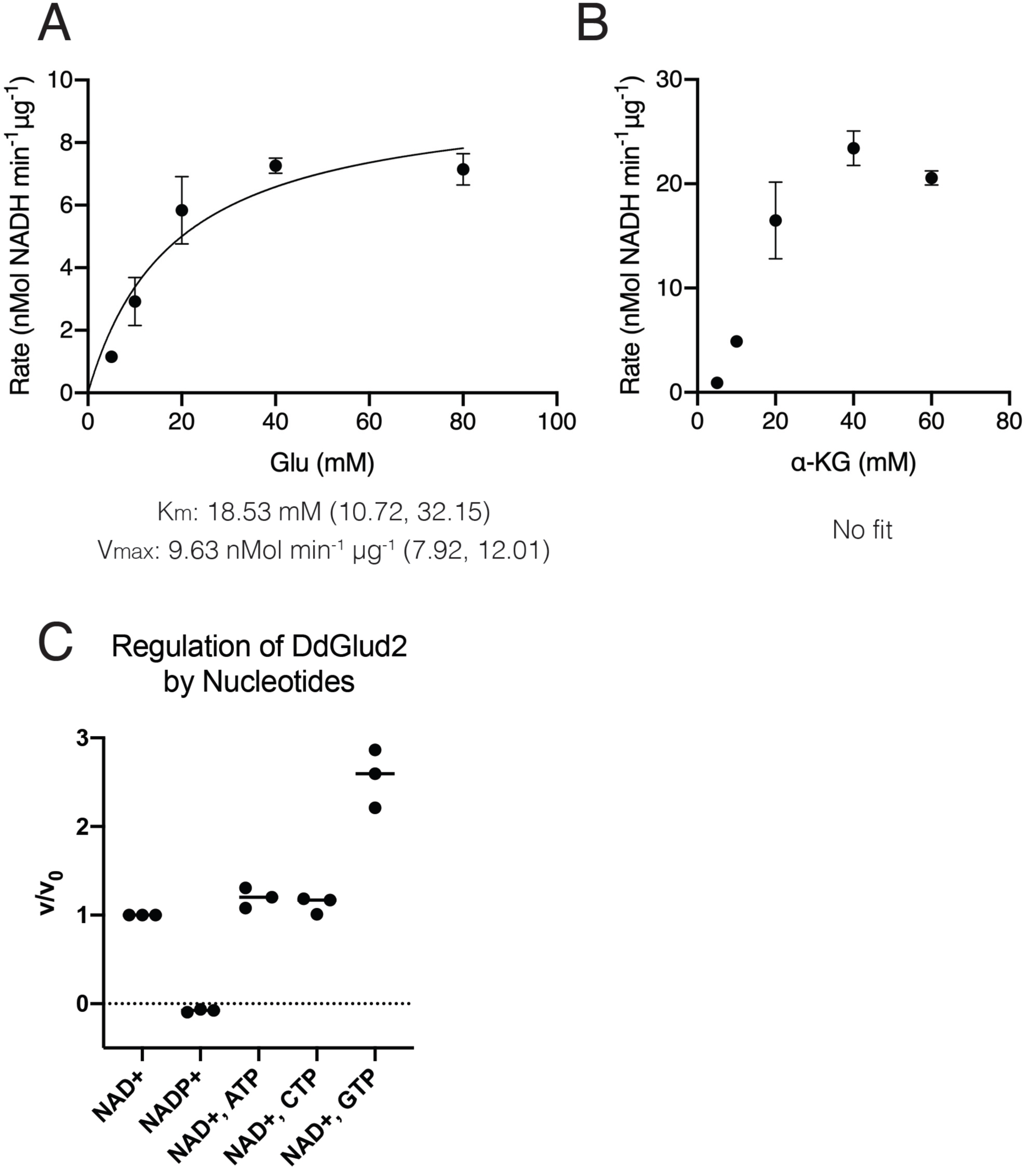
Kinetic properties of *Dd*Glud2. **(A)** Rate plot of *Dd*Glud2 in the presence of excess NAD+ and varying concentrations of glutamic acid. Rate was determined by spectrophotometric measurement of NAD+ reduction to NADH at 340nm. Km and Vmax, calculated by nonlinear regression fitting to Michaelis-Menten kinetics, are indicated below the plot with a 95% confidence interval. **(B)** Rate plot of *Dd*Glud2 in the presence of excess NADH and varying concentrations of α-KG. Rate was determined as in (A). **(C)** Comparison of *Dd*Glud2 reaction rate using NAD+ or NADP as a cofactor, or with 1mM ATP, CTP, and GTP present in the reaction. Reactions in (C) were performed with 10 mM Glu and 1 mM NAD+. The ratio v/v0 expresses the rate of the experimental reaction divided by rate in the presence of NAD+ and no nucleotide (v0.)

**Figure S7.**
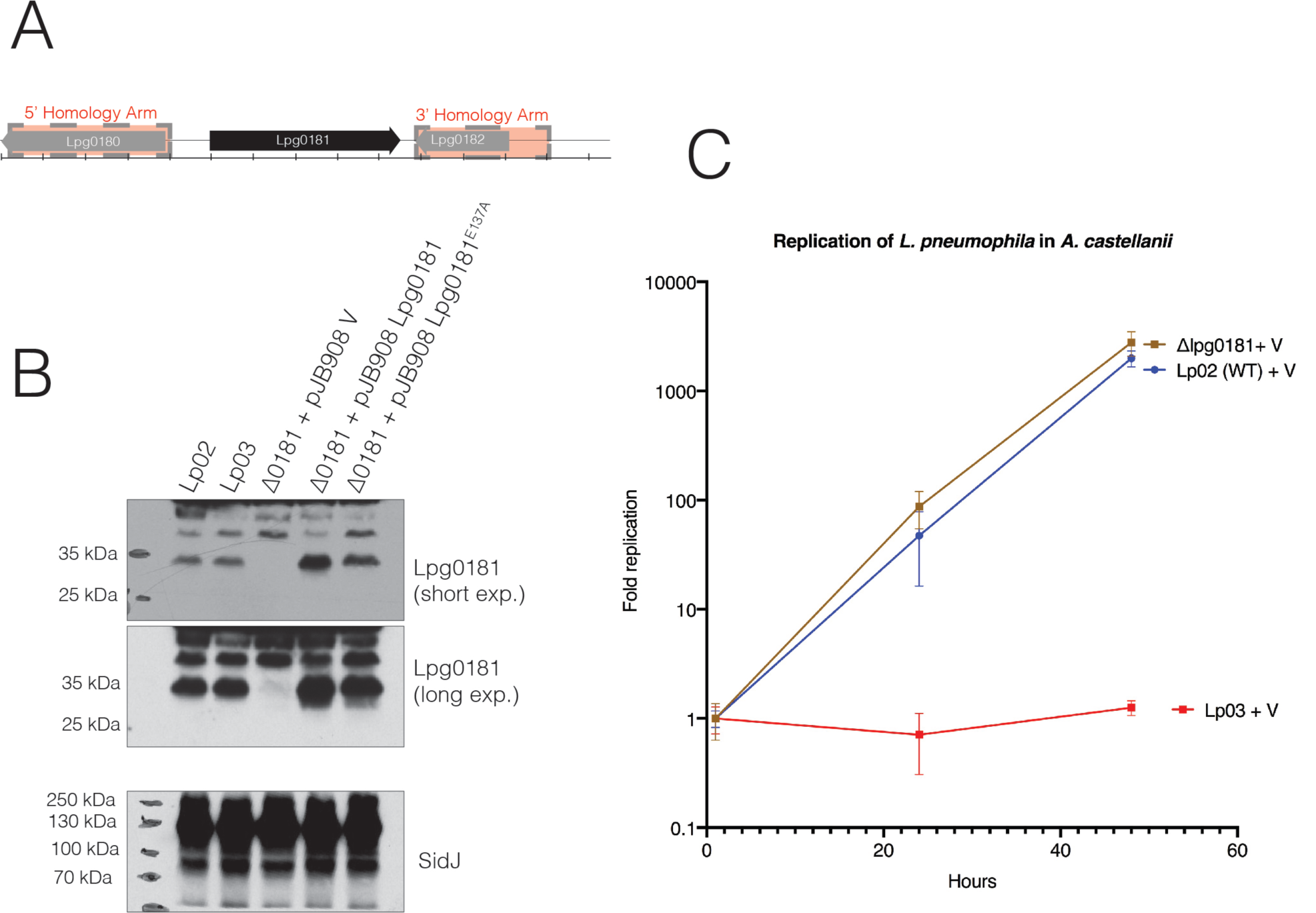
Generation of ΔLpg0181 *Legionella* strains and replication in *A*. *castellani*. **(A)** schematic of the Lpg0181 ORF and the position of the homology arms used to generate the knockout strain. Ticks on the scale bar = 200bp. **(B)** Immunoblotting of *Legionella* bacterial pellets harvested from post-exponential liquid cultures and boiled directly in SDS-PAGE sample buffer. The predicted MW of Lpg0181 is 34.9 kDa. **(C)** Infected amoeba cells were lysed at the indicated time points and bacterial replication was quantified by plating serial dilutions of lysates. Results are obtained from triplicate conditions and are representative of three independent experiments.

**Table S1:** Curated list of proteins enriched from yeast lysate by streptavidin agarose following incubation with Lpg0181, Lpg0181 E137A, and biotin-17-NAD+. Proteins were identified by mass spectrometry. This table includes only proteins unique to the Lpg0181 WT-treated samples. Proteins are listed along with their MASCOT protein score and theoretical molecular weight.

| Top Scoring proteins in Avidin IP - UNIQUE TO WT |  |  |  |
| --- | --- | --- | --- |
| Protein Accession | Name | Score (WT) | Molecular Weight, Da |
| spIP33327IDHE2 | **NAD-specific glutamate dehydrogenase (GDH2)** | 181 | **124,254** |
| spIQ03690ICLU | **Clustered mitochondria protein 1** | 81 | 145,076 |
| spIQ3E764ITMA7 | **Translation machinery-associated protein 7** | 71 | 6,937 |
| spIP31539IHS104 | **Heat shock protein 104 O** | 64 | 101,972 |
| spIP08067IUCRI | **Cytochrome b-c1 complex subunit Rieske, mitochondrial** | 63 | 23,350 |

